# Evaluation of Antibacterial and Antioxidant activity of methanol crude extract of *Aspilia africana* leaves: an in vitro approach

**DOI:** 10.1101/2020.05.27.118745

**Authors:** Adepoju Oluwarinu, Omololu-Aso

## Abstract

*Aspilia africana* (Compositae) is one of such plants considered of great importance in pharmacopeia of traditional medicine. Its leaf is widely used in ethnomedicinal practices in tropical Africa because of its ability to stop bleeding and promote rapid healing of wounds. This study was carried out on the leaf part to determine its antimicrobial and antioxidant potentials of its leaf methanol extract. The methanolic extract of the leaf was subjected to preliminary phytochemical screening and it indicated the presence of saponins, tannin, resin, phlobatannins, and phenols. The *in-vitro* antibacterial test of the methanol crude extract using agar well diffusion method showed broad-spectrum activity with minimum bactericidal concentration of 30, 75 mg/mL for *Klebsiella pneumonia, and Bacillus subtilis* respectively. *In-vitro* antioxidant activities using 2, 2–diphenyl-1-picrylhydrazyl assay indicate that the methanol leaves extract had higher activity than of 92.23 µg/mL compared to standard drugs (Ascorbic acid 1.07mg/mL) and IC_50_ at 4.66. This study concluded that *Aspilia africana* methanol crude extract exhibits dosage-dependent antioxidant potential and could be further explored if it’s are in pure form.

## Introduction

Several thousand plant species have assumed to have contributed to the human diet in the past. They provide food, clothing, shelter, and medicine, and about 150 species have been cultivated for commercial purposes which make plants provide 65% of the world supply of edible protein (Milward, 2005). They are generally divided into cereals, legumes, vegetables, fruits, and nuts, with cereal grains providing almost half (47) of world protein supplies, which makes it a major source of nutrients in developing countries (Milward, 2005). The medicinal uses of plants have been recorded long before history. Much of the medicinal use of plants seems to be developed through observations of wild animals and trial and error (Manuchair, 2000). As time went on each tribe added the medicinal power of herbs in their area to its knowledge base. They methodically collected information on herbs and developed well-defined herbal pharmacopeias (Manuchair, 2000). Many drugs listed as conventional medications were originally derived from plants. Salicylic acid, a precursor of aspirin, was originally derived from white willow bark and the meadowsweet plant (Manuchair, 2000). Cinchona bark is the source of malaria-fighting quinine and other importance drugs (Manuchair, 2000).

World Health Organization survey indicated that about 70 - 80% of the world’s population rely on nonconventional medicine, mainly of herbal sources, in their primary healthcare (Al-Snafi, 2016). This is especially the case in developing countries where the cost of consulting a western style doctor and the price of medication are beyond the capacity of most people (Chan, 2000). Medicinal herbs such as *Aspilia africana* (Compositae) is one of such plants considered of great importance in pharmacopeia. It is a semi-woody herb from a perennial woody rootstock up to 2 meters high. It is very polymorphic and occurs throughout the region on wastelands of the savannah forest. It is also widely distributed across tropical Africa (Dalziel, 1973). *Aspilia africana* is widely used in African folk medicine to stop bleeding, remove corneal opacities, induce delivery, and in the treatment of anemia and various stomachs complain (Iwu, 1993; Adjanohoun *et al*., 1996). *A. africana* is one of the plants that exhibit a wide range of biological activities including antiviral, fungicide and antibacterial activities using various plant parts (Ofusori *et al*., 2008). Phytochemical studies also revealed the presence of saponins and tannins as the most abundant compounds in the plant while flavonoids were the least (Obadoni and Ochuko, 1998). The medicinal plant contained ascorbic acid riboflavin, thiamine, and niacin with a paucity of information on the antioxidant activity of methanol crude extract of the leafy part of *A. africana* (Okwu and Josiah, 2006). Hence this research is to determine the antibacterial and antioxidant potency of the methanol crude extract against selected multiple antibiotic-resistant isolates.

## Materials and method

### Chemical and media

Methanol, 2, 2–diphenyl-1-picrylhydrazyl (DPPH), Peptone, NaCl, Nutrient ager, Mueller Hinton ager. All other chemicals and reagents are of analytical grade.

### Test Organism

The typed clinical test organisms used in this study were collected from the laboratory of the Department of Microbiology, Obafemi Awolowo University. The organisms include Gram-positive *Staphylococcus aureus* (MRSA), *Bacillus stereothemophilus, Bacillus subtilis, and Micrococcus luteus* and Gram-negatives which include *Escherichia coli, Klebsiella pneumonia, and Serratia marcensens, Proteus mirabilis*

### Sample Collection

Uninfected leaves *Aspilia africana* were carefully collected in a disinfected sample collection bag and transported to the university herbarium for further identification. The leaves were rinsed in other to decontaminate and detachment of dirt. The leaves were allowed to air-dry.

### Powdering

After complete drying of the leaves, it was then mechanically ground into fine powdered particles for further analysis

### Preparation of extract

From the powdered air-dried plant sample 300g was weighed and soaked in a solution of 2500ml of 70% methanol for 72 hours at room temperature, it was agitated at intervals of six hours within the 72 hours. The mother liquor was filtered, the filtrate obtained was further concentrated at low temperature (<40°C) under 100 mmHg pressures in a rotary evaporator. The dried extracts were labeled and then stored dry in sterile containers at room temperature until when needed for further procedures.

### Phytochemical Screening (Qualitative Test)

The leave extract was screened for the presence of essential phytochemicals according to the method described by Abulude *et al.* (2001) and Abulude (2007).

### Antibacterial Susceptibility Test

The test isolates were screened against various antibiotics and the results were compared with the result of the test plant part. The already standardized test isolates were be seeded on plated Mueller Hinton agar using spread plate method, followed by placing the antibiotic disc on the surface aseptically and incubated for 18 - 24 hours.

### Preparation of the plant extract

1g (1000mg) of the concentrated plant extract was weighed and dissolved in 10ml of the solvent in, making the stock concentration of the solution (plant extract) 100mg/ml. double fold dilution of plant extract was prepared by pipetting 5ml in to test tube containing 5ml of the diluent to make 50mg/ml, the same step is repeated for 25mg/ml,12.5mg/ml,6.25mg/ml, 3.252mg/ml and 1.625mg/ml respectively.

### Extract susceptibility test

Test bacteria isolates were seeded on the surface of already prepared Mueller Hinton, 6mm well was bored in the agar using cork borer and calibrated micropipette was used to transfer 0.1ml of the extracts of the prepared concentrations into the already labeled well. Streptomycin was used as negative control. The antibacterial agent was allowed to diffuse into the agar for about 30 minutes before incubation at 37°C for 18 – 24 hrs. The sensitivity, minimum inhibitory concentration (MIC) and minimum bactericidal concentration (MBC) of the plant extract was evaluated by measuring the diameter of the zone of inhibition for each of the plate using a transparent plastic ruler and the mean was being taken.

### Antioxidant Test

The radical scavenging ability of the oil was determined using the stable radical DPPH (2, 2-diphenyl-1-picrylhydrazyl hydrate) as described by Brand-Wasiams *et al.* (1995). The reaction of DPPH with an antioxidant compound which can donate hydrogen leads to its reduction (Blois, 1958).The change in colour from deep violet to light yellow was measured spectrophotometrically at 517nm.

### Statistical Analysis

Data generated was analysed using appropriate statistical software, presented diagrammatically in form of pie chart and bar chart where necessary.

## RESULT AND DISCUSSION

The preliminary phytochemical screening (Table 1) of methanolic crude extract showed high presence of tannins, saponin, resin, phlobotanins, and phenol while other typical plant chemicals namely glycosides, sterols, flavonoids, and carbohydrates (reducing sugars) were absent. This was similar to the study carried out by Johnson *et al*. (2011) and Oko and Agiang (2011). The observed result can due to varying extraction solvent, climatic condition across geographical locations and plant age. The extent of the inhibitory activity of plant against microorganisms is a complex relationship which is determined by kind and concentration of the bioactive substances present. These bioactive substances are usually responsible for the pharmacological activities of medicinal plants as reported by El-Tantaway *et al*. (1999). The antibacterial activity of *A. africana* was observed to be broad spectrum as shown in table 2 against nine (9) multidrug resistant bacteria test isolates. The minimum inhibitory concentration (MIC) of the crude extract on Gram-negative bacteria were exhibited at concentration between 12.5 -100 mg/ml and Gram-positive between 25 – 100 mg/ml (table 2). The observed result of the antibacterial activity was in line with previous study by Adetunji *et al*. (2012). Furthermore, MBC observed against test strains was range between 30 – 200 mg/ml, while *Klebsiella pneumonia* was killed at the lowest concentration of 30 mg/ml and *B. stereothermophilus* and *Staphylococcus aureus* at 200 mg/ml (table 2) which indicates that crude extract actively inhibited the Gram-negative bacteria at a lower concentration compare to the Gram positive bacteria which is in contrary to study by Johnson *et al*. (2011). This can be suggestive that the active antimicrobial agent is concentration, microbial population and method dependent or can evade the peptidoglycan layer barrier of the Gram negative by altering the integrity of the cell wall as to compare to Gram positive cell wall.

**Table 1:**
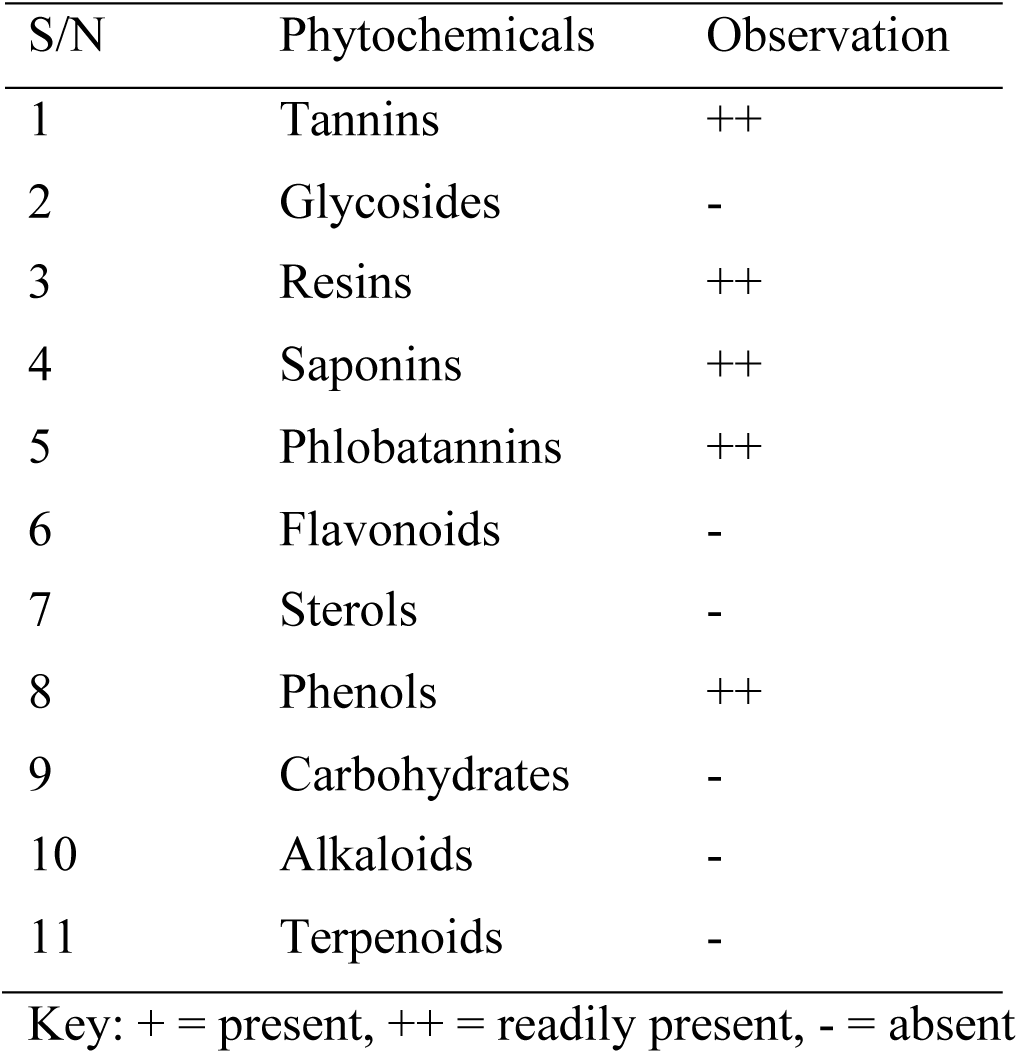
Phytochemical screening result for methanol extract

**Table 2:** Antibacterial sensitivity test result of Methanol crude extract

| S/N | CODE | Positive control | MIC (mg/ml) | MBC (mg/ml) |
| --- | --- | --- | --- | --- |
| 1 | *Serratia marcescens* | - | 12.5 | 75 |
| 2 | *Klebsiella pneumonia* | - | ND | 30 |
| 3 | *Micrococcus luteus* | - | 50 | 75 |
| 4 | *Escherichia. coli* | - | 100 | ND |
| 5 | *B. stereothermophilus* | - | 100 | 200 |
| 6 | *Bacillus subtilis* | - | 25 | 75 |
| 7 | *S. aureus* | - | 100 | 200 |
| 8 | *Proteus mirabilis* | - | ND | 200 |
| 9 | *S. aureus (MRSA)* | - | 25 | ND |

Key ND: Not determined

Antioxidants are tremendously important substances which posses the ability to protect the body against free radical induced oxidative stress (Badakhshan *et al*., 2012). 2, 2–diphenyl-1-picrylhydrazyl (DPPH) radical scavenging activities of the methanol extract of *A. africana* were shown in table 3 and figure 1.The *A. africana* leaves antioxidant activity was observed to significantly increase with decreasing concentration of the crude extract. The antioxidant activity on DPPH was believed to be as a result abundance of phenolic biocompound. Plant phenolic biocompound acts as reducing agent and antioxidant by donating its hydroxyl group. The scavenging abilities of the extracts were significantly higher than ascorbic acid (standard drug) at a lower concentration (table 3), it was evident that the MER possesses potential to act as a primary antioxidant. The quality of the antioxidants in the extracts was determined to double inhibition concentration (IC 50) value of the standard drug as shown in table 3.

**Table 3:** Percentage (%) free radical scavenging inhibition of MER.

| S/N | Sample Concentration (mg/ml) | Ascorbic acid (% Scavenging inhibition) | *Aspilia africana* (% Scavenging Inhibition) |
| --- | --- | --- | --- |
| 1 | 10 | 59.96 | 58.96 |
| 2 | 5 | 35.43 | 50.68 |
| 3 | 2.5 | 19.54 | 57.16 |
| 4 | 1.25 | 4.96 | 93.53 |
| 5 | 0.0625 | 3.17 | 89.17 |
| 6 | 0.03125 | 1.07 | 92.23 |
| 7 | IC<sub>50</sub> | 8.08 | 4.66 |

**Figure 1:**
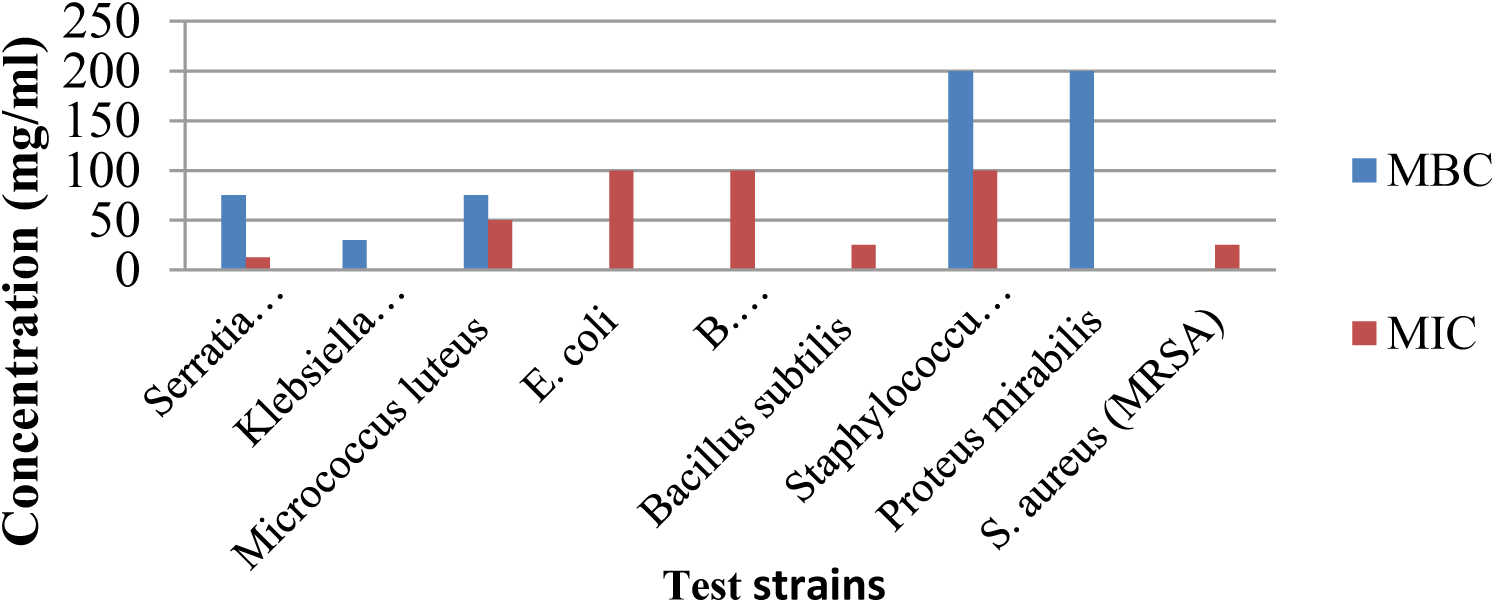
Showing the minimum inhibitory concentration (MIC) and Minimum bactericidal concentration….

**Figure 2:**
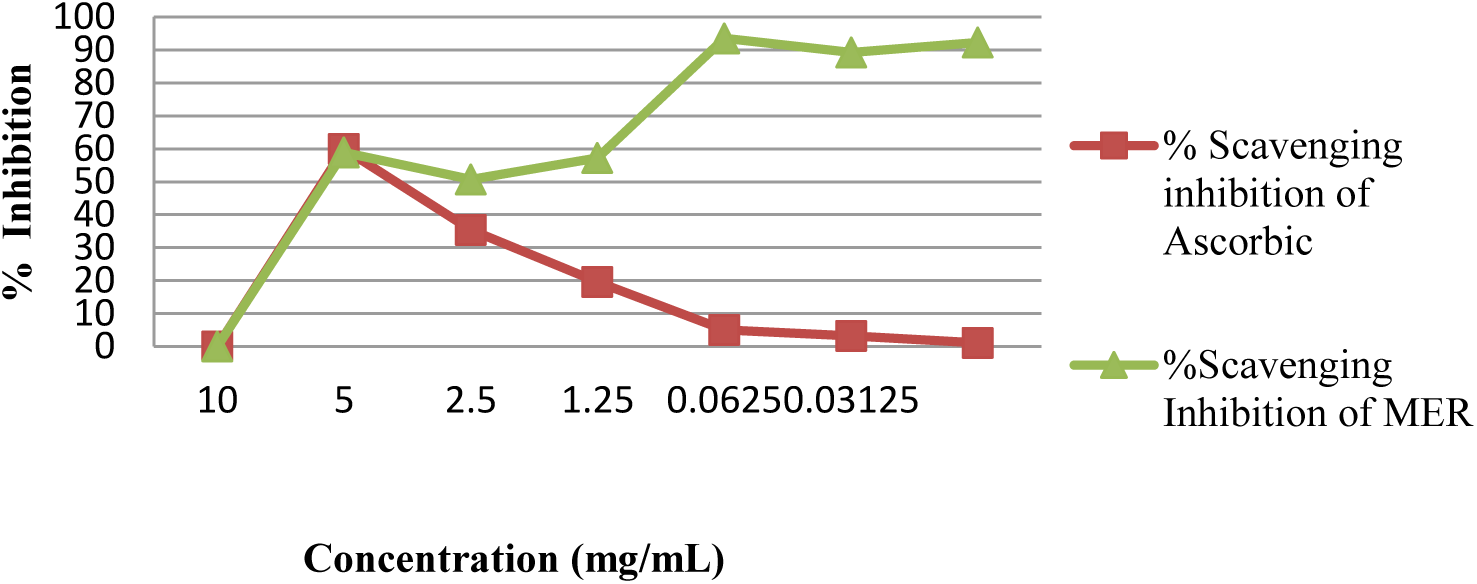
Showing the percentage scavenging inhibition of Ascorbic acid and methanol crude.

## CONCLUSION

From this study, the methanolic crude extract of *Aspilia africana* has been reaffirmed to possess a broad spectrum of antibacterial activity but with better activity against Gram negative bacteria. The total phenolic contents of the MER contribute to the antibiotic and antioxidant property of the plant. Therefore the extraction of active compounds from the plants leave for qualitative and quantitative analysis could be an alternative cheaper natural therapeutic drug compare to available synthetic drugs.

